# Thrombo-inflammatory endothelial signatures in *JAK2*-mutated myeloproliferative neoplasms

**DOI:** 10.64898/2026.02.16.706250

**Authors:** Salma A.S. Abosabie, Alexandra Boye-Doe, Mohammed Ali, Nikolai Podoltsev, David Stegner, Lourdes M. Mendez, Anish V. Sharda

## Abstract

**Background:** Classical myeloproliferative neoplasms (MPN)—essential thrombocythemia, polycythemia vera, and primary myelofibrosis—are characterized by clonal hematopoiesis, overproduction of mature blood cells, and a high burden of thromboembolic events. Although thrombosis is the leading cause of morbidity and mortality in MPN, the contribution of the vascular endothelium remains incompletely defined. We investigated patient-derived endothelial colony-forming cells (ECFCs) as a surrogate for vascular endothelium in individuals with JAK2 V617F–mutated MPN.

**Methods:** ECFCs were cultured from peripheral blood of patients with MPN and healthy controls, phenotyped for thrombo-inflammatory and adhesive markers, tested for JAK2 V617F, and profiled by bulk RNA sequencing. Functional assays assessed endothelial-dependent factor Xa generation. Transcriptomes were benchmarked against public HUVEC reference datasets processed through an identical quantification pipeline.

**Results:** ECFCs were obtained more frequently and in greater numbers from patients with MPN than from controls, indicating enhanced endothelial regenerative or activation potential. MPN ECFCs exhibited increased von Willebrand factor and P-selectin expression and release, along with elevated endothelial cell–dependent factor Xa generation, consistent with a thrombo-inflammatory, procoagulant phenotype. JAK2 V617F was not detected in any ECFC colonies, supporting a non-clonal origin of these endothelial abnormalities. Transcriptomic analysis identified 289 differentially expressed genes in MPN versus control ECFCs, with pathway enrichment revealing coordinated dysregulation of blood coagulation, platelet activation, plasminogen regulation, vascular permeability, extracellular matrix organization, and angiogenesis. Benchmarking against HUVEC datasets confirmed strong endothelial identity of ECFC-derived cells, with MPN-associated changes reflecting endothelial activation rather than loss of endothelialness.

**Conclusions:** ECFCs from patients with JAK2-mutated MPN display functional and transcriptomic signatures of endothelial dysfunction in the absence of detectable driver mutations. These findings support a model in which a primed, thrombo-inflammatory endothelium cooperates with clonal hematopoiesis to promote the heightened thrombotic risk characteristic of MPN.

## Introduction

Myeloproliferative neoplasms are bone marrow disorders characterized by excess clonal hematopoiesis resulting in elevated peripheral blood counts.^1^ Among several distinct clinicopathologic entities defined by the World Health Organization classification system of tumors of the hematopoietic and lymphoid tissues, polycythemia vera (PV) and essential thrombocythemia (ET) are the most common.^2^ Somatic mutations in Janus kinase 2 (*JAK2*), particularly *V617F* mutation in exon 14 of *JAK2,* are the most frequent genetic alterations associated with MPNs, found in >98% of all cases of PV and ∼50% of ET.^1^ Somatic mutations in *CALR* and *MPL* are the next more frequent genetic alterations, found exclusively in ET or primary myelofibrosis.^1^

One of the hallmarks of MPNs, particularly PV and ET, is a substantially elevated risk of thromboembolic events (TEs).^3,4^ The risk of TE is about 5-7-fold elevated in MPNs, and can involve either arterial or venous circulations.^5^ TEs, frequently idiopathic and involving unusual sites such as splanchnic venous system, often lead to the diagnosis of MPNs and can be associated with significant morbidity and mortality.^6^ Although the risk of TE in MPN, particularly PV and ET, is well recognized, it’s pathophysiology remains poorly understood. Both cellular and humoral factors, including platelet hyperreactivity, neutrophil and monocyte activation, hyper-viscosity, procoagulant extracellular vesicles, cytokines and coagulation abnormalities, have all been reported.^6–10^

Endothelial cells, which line the blood vessels and maintain integrity of the circulatory system, play a critical role in pathogenesis of thrombosis. Endothelial dysfunction has also been in MPN, however the exact underlying mechanisms remain incompletely understood.^7,11–13^ Elevated levels of soluble endothelial-derived factors such as P- and E-selectin, von Willebrand factor (vWF), thrombomodulin and protein-disulfide isomerase has been reported in MPN.^7,14–16^ Additionally, elevated plasma angiogenic factors such as vascular endothelial growth factor (VEGF) has been shown.^17^ However, exact underlying mechanisms remain incompletely understood and the contribution of endothelial dysfunction remains underexplored. Also, in dispute is whether MPN endothelium could possess driver mutation such as *JAK2 V617F*. This is mainly due to difficulty in retrieving endothelial cells from the lining of the blood vessels from living subjects. To address this, endothelial colony forming cells (ECFCs), are being increasingly utilized. ECFCs are progeny of circulating and transplantable myeloid-derived endothelial progenitor cells that can be easily harvested from small volume of peripheral blood.^18^ Although, the origin of ECFCs is not entirely clear but these cells are both phenotypically and genotypically bonafide endothelial in nature, and have significantly improved understanding of disorders such as early-onset atherosclerosis, thromboembolism of sickle cell disease, von Willebrand disease, and rheumatologic disorders, among others.^19–25^ ECFCs have been examined previously in MPN, but prior work has been largely focused on descriptive or functional phenotypes. In this study, we performed an integrated phenotypic and transcriptomic analysis of endothelial colony-forming cells (ECFCs) derived from a cohort of patients with *JAK2 V617F*–mutated MPN. Our data reveal marked endothelial dysfunction in MPN ECFCs, characterized by impaired vascular phenotype and widespread dysregulation of endothelial gene expression programs.

## Materials and methods

### Study approval, patients and blood collection

The protocol was approved by the Institutional Review Board at Yale New Haven Hospital. Studies were conducted in accordance with the Declaration of Helsinki. Written informed consent was obtained from all study participants before inclusion in the study and blood draw. Table S1 summarizes the inclusion exclusion criteria. MPN patients were eligible if they were 18 years of age or older and had a known diagnosis of *JAK2-*mutated MPN. Controls were healthy laboratory volunteers, considered to be in good health, specifically without any known disorders of hemostasis or thrombosis, cardiac disease, or cancer. A total of 7 MPN and 13 control subjects were enrolled. A total of 30-36 ml of heparinized peripheral whole blood was collected from each subject at enrollment.

### Reagents

All reagents, antibodies and commercial kits used are presented in major resources table (**Table S2)**.

### ECFC culture

ECFC were cultivated as previously described.^18^ Briefly, PBMC fraction of whole peripheral blood from each subject was plated on two collagen-coated 48-well plates in endothelial cell media (EBM-2;Lonza) supplemented with endothelial bullet kit (Lonza) and 20% FBS (complete media). Plates were monitored closely for colonies for up to 6 weeks and media changed 2-3 times per week. Cells growing in a cobblestone pattern were passaged to larger cultures once 70-80% confluent, and classified as ECFCs if: 1) negative for surface CD45 2) positive for surface PECAM1 3) and positive for vWF (**Table S1**).

### Flow Cytometry

ECFCs were detached using accutase, washed with PBS, and incubated with fluorophore-labeled antibodies or isotype controls in FACS buffer for 30 minutes on ice. Data was then acquired on BD LSR Fortessa flow cytometer and analyzed using BD FlowJo Software.

### vWF ELISA

VWF enzyme-linked immunosorbent assay (ELISA) was performed as previously described.^26^

### Immunofluorescence microscopy

Immunofluorescence microscopy was performed as previously described.^27^ Cells were cultured on chamber slides, washed with PBS, fixed in 4% paraformaldehyde, permeabilized with 0.1% triton-X, and incubated with rabbit anti-vWF antibody followed by Alexa Fluor 488 anti-rabbit secondary. DAPI was used for counterstaining and immunofluorescence images captured on an inverted TCS SP8 Leica Microscope.

### MTT assay

3-[4,5-dimethylthiazol-2-yl]-2,5 diphenyl tetrazolium bromide (MTT) proliferation assay was performed per manufacturer’s protocol.

### DNA and RNA isolation

RNA and DNA were extracted using Qiagen kits per the manufacturer’s protocol.

### *JAK2 V617F* mutation analysis

***JAK2 V617F*** mutation analysis was carried out using predesigned qPCR primers (Invitrogen) according to manufacturer’s protocol. The following primer set was used: 1) JAK2 V617F primer assay (Assay ID Hs00000940_mu Gene JAK2) JAK2_12600_mu; 2) JAK2 reference gene primer assay (Assay ID H500001020_rF Gene JAK2).

### Bulk RNA sequencing and data analysis

Bulk RNA sequencing was performed on single colonies using Illumina Novaseq S4 by the Keck Biotechnology Resource Laboratory at Yale. The sequencing reads for each of the samples are aligned to the GRCh38 human reference using HISAT2 (Kim, et.al., 2015). Gene-level read counts are generated using StringTie2 and its prepDE.py tool (Kovaka, et.al., 2019), based on annotations from the ENCODE v27 GTF file. Sample and experiment quality metrics are generated using Picard (Picard Toolkit, 2019), and TPM counts are generated using Ballgown (Pertea, et.al., 2016). Differential gene expression is performed using DESeq2 (Love, et.al., 2014), and the DESeq2 analysis results are submitted to the Ingenuity Pathway Analysis software (QIAGEN) for a Core Analysis to report pathway enrichment. Pathway enrichment analysis was performed using the Enrichr suit (Ma’ayan lab) ^28–30^ and Metascape gene analysis tool ^31^.

### HUVEC-reference transcriptomic analysis

Bulk RNA-seq from ECFCs (5 controls, 10 MPN) was quantified at the transcript level using Salmon against the GENCODE GRCh38 transcriptome and summarized to gene-level counts with tximport. Gene counts were converted to log2(CPM+1) for downstream comparisons. Public HUVEC RNA-seq FASTQs were downloaded from ENCODE (experiment accession ENCSR000EYS), quantified using the same Salmon/GENCODE pipeline, and used as a reference. HUVEC-likeness was defined as the Pearson correlation between each ECFC gene-expression profile and the HUVEC reference profile. In parallel, ECFCs and HUVECs were jointly embedded by principal component analysis (PCA), and distance to HUVEC was computed as Euclidean distance from each ECFC to the HUVEC centroid in PCA space. An endothelial identity score and additional program scores (NF-κB feedback, TNF/NF-κB activation, IFN/STAT1) were computed per sample as the mean z-scored expression of curated gene sets. Group differences (Control vs MPN) were assessed using two-sided Wilcoxon rank-sum tests with rank-biserial effect sizes; figures display individual samples with median/IQR and mean±SD overlays.

### Statistics

Data analysis and graphing were performed using GraphPad Prism software. Continuous variables are presented as mean ± SD unless otherwise specified. For comparisons between two independent groups, unpaired two-tailed Student’s t-tests were used when data met assumptions of normality and equal variance; when these assumptions were not satisfied, the non-parametric Mann–Whitney U test was applied. Categorical variables (e.g., proportion of samples yielding at least one ECFC colony) were compared using Fisher’s exact test. A P value <0.05 was considered statistically significant. For transcriptomic datasets, differential expression and pathway enrichment analyses were performed using standard RNA-seq packages as detailed above, with multiple-testing correction by the Benjamini–Hochberg method; genes or pathways with false discovery rate (q value) <0.05 were considered significantly enriched. Differential expression between Control and MPN BOECs was evaluated at the gene level using log2(CPM+1) values, with a priori focus on canonical endothelial markers and inflammatory/thrombo-inflammatory genes; group differences were tested using two-sided Wilcoxon rank-sum tests with rank-biserial effect sizes.

## Results

### Clinical characteristics of patients with MPN

The clinical characteristics of patients with MPN are highlighted in Table 1. Age range was between 45-89 years and 4 were females. The primary diagnosis was ET in 3 patients, PV in 3 patients and PMF in 1. All patients had *JAK2 V617*-mutated MPN, with variable allele frequency ranging from 4.5%-57.5%. Three patients with MPN were on low-dose aspirin, and 4 on anticoagulant medication. Four patients were receiving cytoreductive therapy with hydroxyurea, while 2 were on JAK1/2-inhibitor ruxolitinib. In addition, 13 healthy volunteers with no significant medical or medication history were enrolled as controls. ECFCs were successfully harvested from 6 out of 7 total patients with MPN vs. 5 out of 13 controls (**Figure S1**).

**Table 1.** Clinical characteristics of patients with MPN (VAF, variable allele frequency; AC, anticoagulant; ET, essential thrombocythemia; PV, polycythemia vera; PMF, primary myelofibrosis; ASA, aspirin; DVT, deep vein thrombosis; PE, pulmonary embolism; CVA, cerebrovascular acciden7

| Patient | Age (Years) | Gender | Primary Diagnosis | Time Since Diagnosis (Years) | JAK2 V617F VAF (%) | History of Thrombosis | AC | ASA | Current Treatment | Thrombotic events |
| --- | --- | --- | --- | --- | --- | --- | --- | --- | --- | --- |
| MPN 1 | 72 | F | ET | 5 | 11% | Yes | Yes | No | Hydroxyurea<br>Apixaban | Splanchnic venous thrombosis |
| MPN 3 | 69 | F | PV | 1 | 6.5 | Yes | Yes | No | Apixaban<br>Ruxolitinib | Arterial thrombosis;<br>CVA |
| MPN 4 | 66 | M | ET | 6 | 19% | Yes | No | No | ASA<br>Hydroxyurea<br>Plavix | CVA |
| MPN 5 | 45 | F | ET | 9 | N/A | Yes | Yes | No | Hydroxyurea<br>Rivaroxaban<br>Therapeutic phlebotomy | Budd-Chiari Syndrome;<br>PE |
| MPN 6 | 67 | M | PV | 6 | N/A | Yes | No | Yes | ASA<br>Hydroxyurea<br>Therapeutic phlebotomy | Recurrent superficial vein thrombosis |
| MPN 7 | 85 | M | PV | 1 | 57.5 | No | No | Yes | ASA<br>Ruxolitinib<br>Therapeutic phlebotomy | None |
| MPN 8 | 89 | F | PMF | 5 | 4.5 | Yes | Yes | No | Hydroxyurea<br>Rivaroxaban<br>Therapeutic phlebotomy | DVT/PE |

### ECFC culture and characterization

Peripheral Blood Mononuclear Cells (PBMCs) isolated from whole blood samples were plated on collagen-coated 48-well plates in complete endothelial growth media and followed for appearance of ECFC colonies. Table S1 lists the criteria for calling harvested colony forming units as ECFCs. ECFCs possessed the characteristic cobblestone-like morphology of endothelial cells, were positive for surface expression of endothelial marker (CD144) and negative for surface expression of leukocyte common antigen CD45 (**Figure 1A-C**). Additionally, all ECFCs expressed vWF in characteristic cigar-shaped Wiebel-Palade bodies on immunofluorescence microscopy (**Figure 1D**).

**Figure 1:**
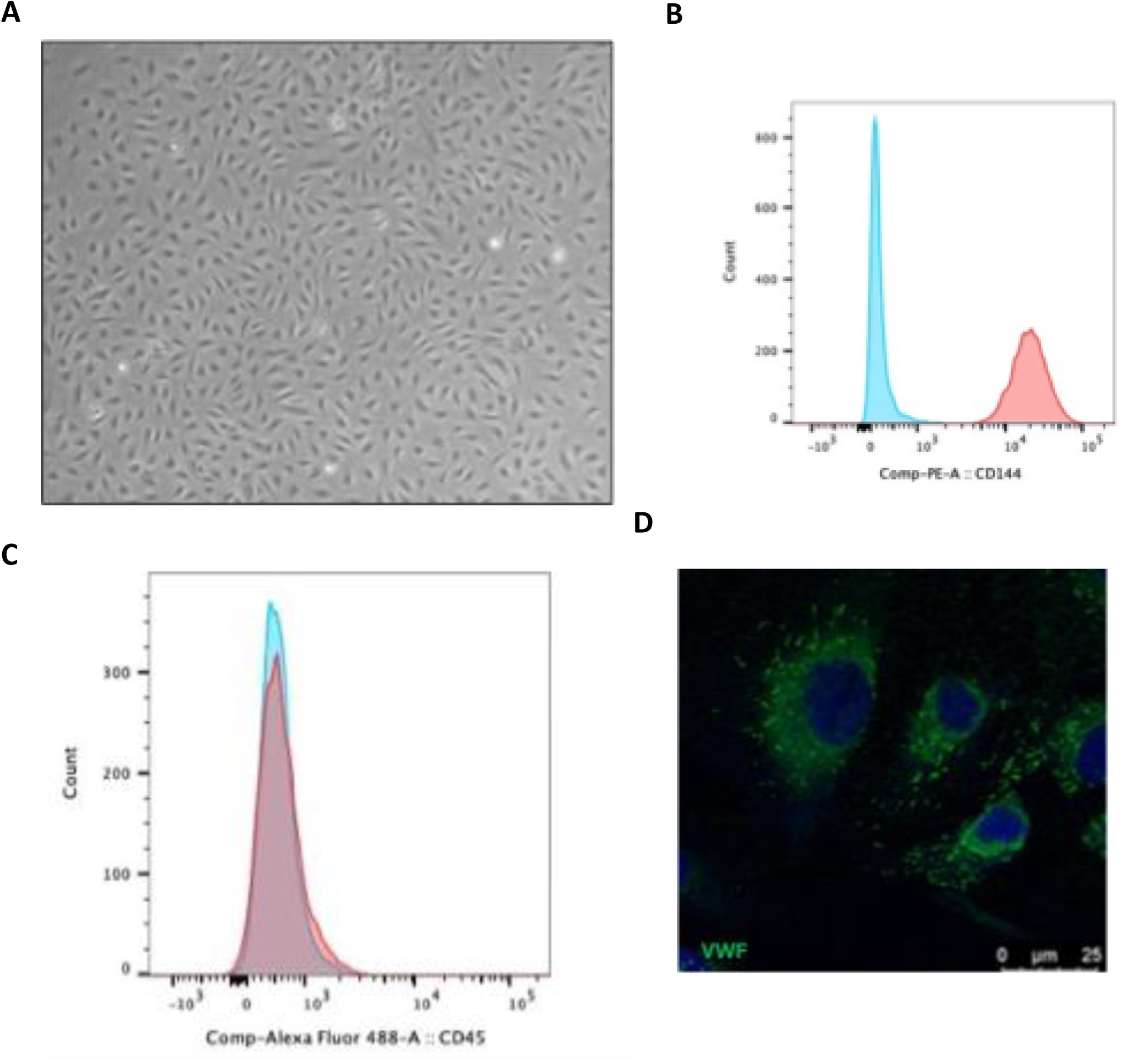
Characterization of endothelial colony forming cells. (A) Brightfield microscopy of a ECFC colony showing a typical monolayer growth and cobblestone appearance characteristic of primary endothelial cells in culture (B-C) ECFCs in culture were detached, washed, and then incubated with PE-labeled anti-CD144 (PECAM-1) antibody (B) or Alexa Fluor 488-labaled anti-CD45 antibody (C) and surface expression of these markers determined by flowcytometry (blue, isotype control; red, CD144 or CD45). D) Immunofluorescence micrograph showing presence of vWF-containing (green) cigar-shaped Weibel Palade bodies in ECFCs (blue, DAPI; scale bar 25 micron).

Whether MPN endothelium can also acquire somatic *JAK2 V617F* mutation remains disputed. We, therefore, tested ECFCs from our cohort *for JAK2 V617F* mutation. DNA was extracted from all ECFC colonies obtained and *JAK2 V617F* mutation analyzed using qPCR (limit of detection VAF < 0.01%) (**Figure S2**). None of the ECFC colonies from our cohort were found to be positive for mutant *JAK2*. As expected, all controls colonies were also negative for *JAK2 V617F* mutation.

### Elevated ECFC regenerative capacity in MPN

ECFCs were successfully harvested in 6 out of 8 patients with MPN, as compared with 5 out of 13 controls (**Figure S1**). The zero-colony outcome, defined as the absence of any ECFC colonies after 6 weeks of PBMC culture, was more frequent in controls as compared with the MPN cohort (54% vs. 25%) (**Figure 2A**). Moreover, the mean number of ECFC colonies isolated from each study subject was significantly higher in the MPN cohort compared with controls (7 vs. 1.3, p=0.0033) (**Figure 2B**).

**Figure 2.**
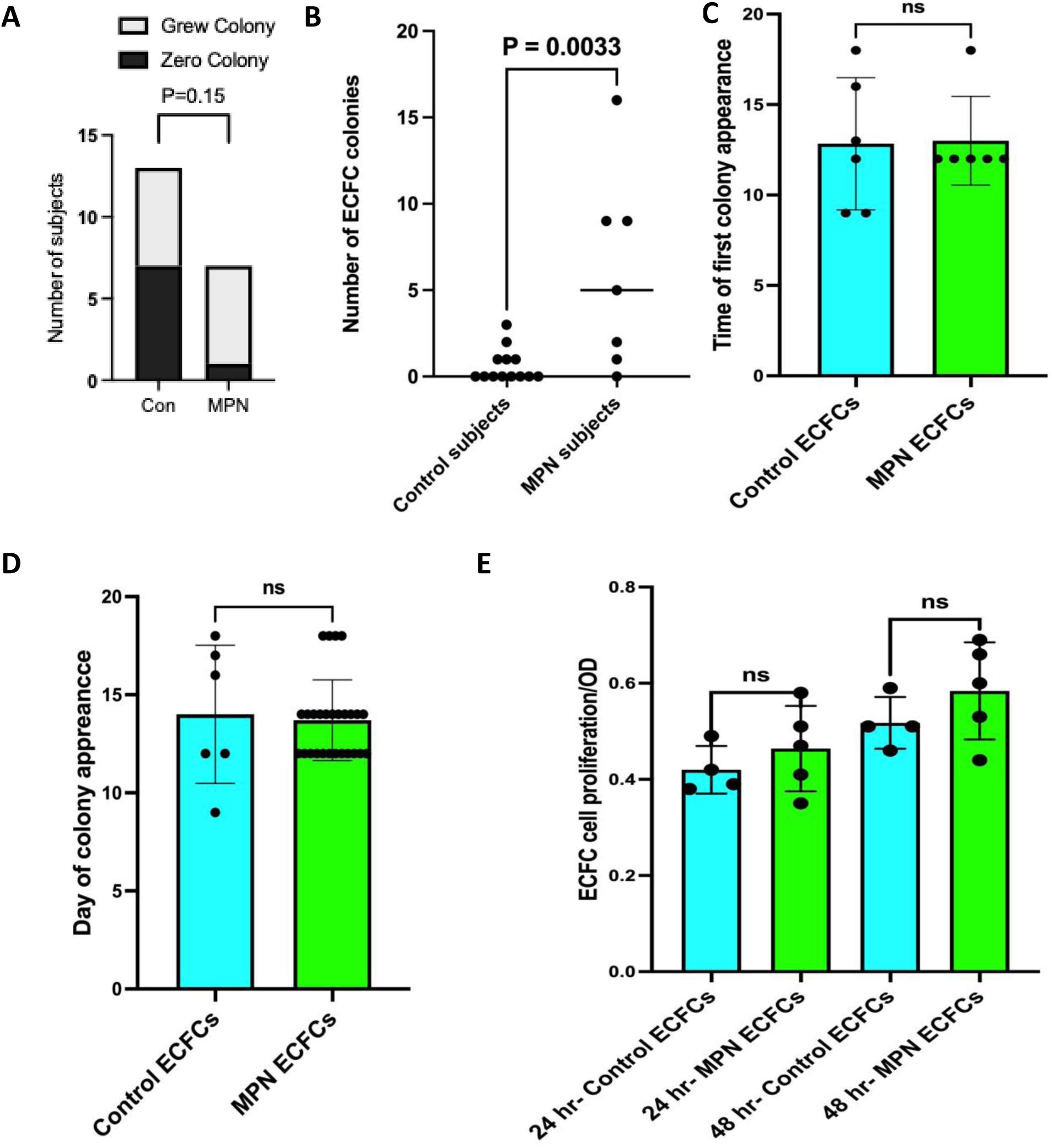
Elevated endothelial regenerative capacity in MPN. (A) Zero-colony outcomes (subjects with no ECFC colonies) are shown as proportions; p-value were computed by Fisher’s exact test (B) number of ECFC colonies harvested per subject comparing MPN vs controls is shown. Mean is depicted by the horizontal line; p-value computed by Wilcoxon rank-sum (C-E) Bar graphs showing time to first ECFC colony appearance (C) time to appearance of individual ECFC colony (D) and cell proliferation rate as determined by the MTT assay at 24 and 48 hours (E). Individual values with mean with SD are shown.

While patients with MPN had a higher mean number of ECFC colonies, the day of colony appearance (13.7 vs. 14 days; p=0.4 in control and MPN ECFCs respectively) and the time to first colony appearance (12.8 vs. 13.0 days; p=0.40) were similar in both cohorts (**Figure 2C-D**). The day of colony appearance was recorded for each subject to track the timing of endothelial colony formation, and day 0 was defined as the day PBMCs were seeded into culture plates. This was supported by the cell proliferation rates at 24 hours estimated by the MTT assay (OD 0.5 vs. 0.65; p=0.15), which did not differ between MPN and controls (**Figure 2E**). This likely suggests that there is an increased endothelial regenerative capacity in MPN, while the proliferation rate of MPN endothelium remains unaltered as compared with controls.

### MPN ECFCs are characterized by a thrombo-inflammatory phenotype

Next, we evaluated the expression of vWF and P-selectin, two markers of thrombo-inflammation, in MPN ECFCs. vWF expression was determined on ECFC lysates using an ELISA. As compared with controls, MPN ECFCs demonstrated a significantly increased vWF content (5260 vs. 1160 arbitrary units per mg of lysate; p=0.004) (**Figure 3A**). Additionally, MPN ECFCs were characterized by augmented vWF release as compared with controls (**Figure 3B**). Similarly, P-selectin surface expression as determined by flow cytometry was significantly elevated in MPN ECFCs compared with controls (mean MFI 358 vs. 190; p=0.008) (**Figure 3C**). Notably, there was intra-subject heterogeneity in vWF and P-selectin expression observed across colonies within the MPN cohort, but a strong correlation existed between vWF and P-selectin surface expression (R = 0.74). (**Figure 3D; Figure S3**). These data support increased endothelial-dependent thrombo-inflammation in MPN.

**Figure 3:**
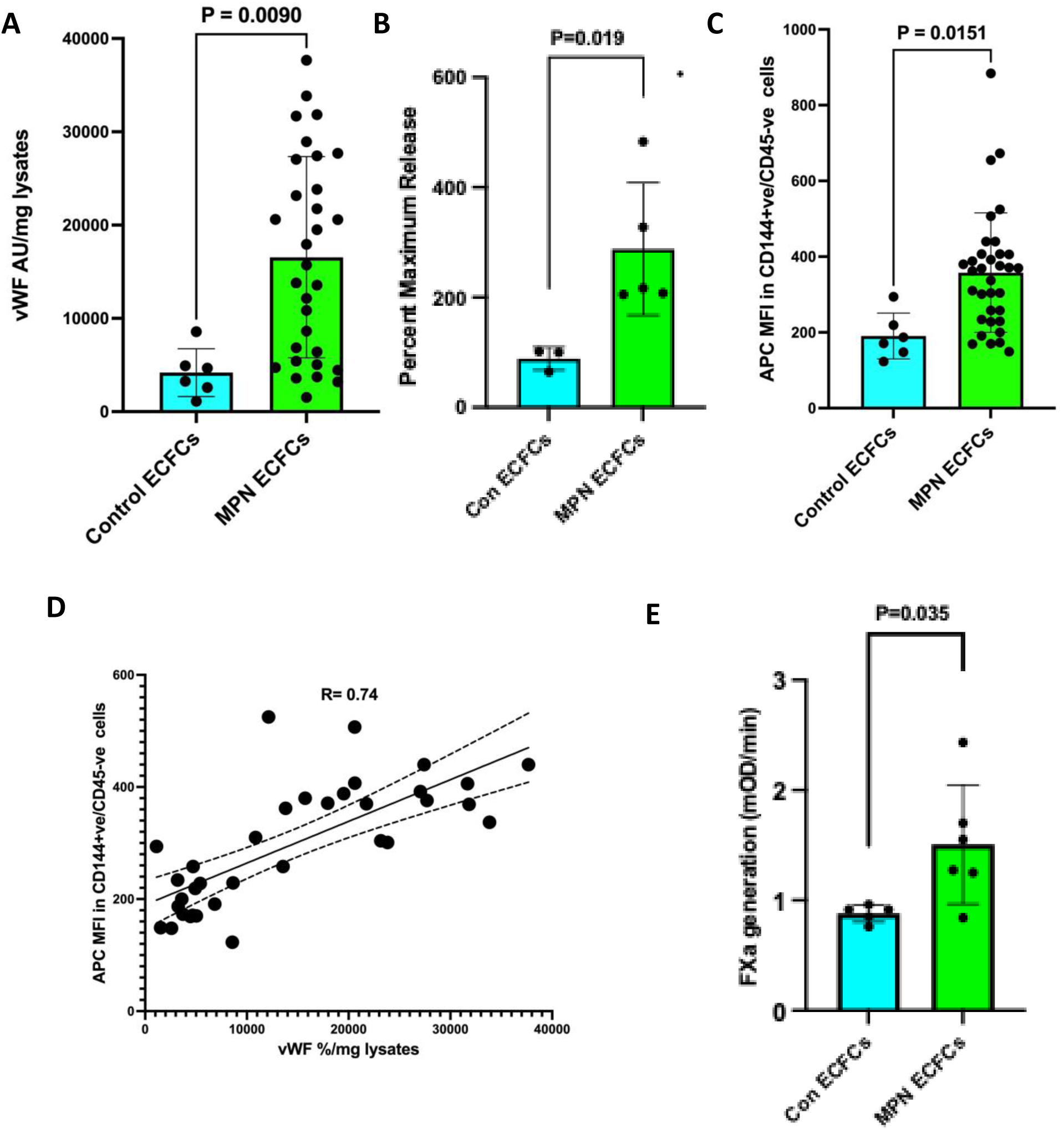
MPN endothelium is characterized by a thrombo-inflammatory phenotype. (A) Scatter and bar plots illustrating vWF antigen per mg of lysate (arbitrary units (AU)) of individual ECFC colony determined by ELISA. Individual values with mean and SD are shown. p-value were computed by Welch’s t-test (B) Scatter and bar plots showing vWF antigen in culture media of ECFC monolayers (one colony per subject), as determined by ELISA, 30 minutes after of stimulation with 100 µM histamine. One colony from each subject was tested. Individual values with mean and SD are shown; p-value were computed by Welch’s t-test (C) ECFC monolayers were detached, washed, and then incubated with APC-labeled anti-P-selectin antibody. P-selectin surface expression was determined by flowcytometry. Scatter and bar plots showing P-selectin expression of individual ECFC colonies together with mean and SD, comparing MPN and controls; p-value were computed by Welch’s t-test (D) Correlation between vWF content and P-selectin surface expression in every ECFC colony in MPN cohort. (E) Scatter and bar plots showing rate of factor Xa generation (mOD/min) in monolayers of ECFCs stimulated with 100 ng/ml LPS for 4 hours. Factor Xa generation determined by rate of cleavage of chromogenic factor Xa substrate, after addition of chromogenic substrate together with coagulation factors VIIa and X to cells. One colony from each subject was tested. Individual values are shown together with mean and SD; p-value were computed by Welch’s t-test.

### Increased endothelial procoagulant activity in MPN

Next, we evaluated if MPN endothelium could also support increased fibrin generation *in vitro*. For this we employed a factor Xa generation assay adapted to be endothelial-cell dependent, as previously described.^32^ One ECFC colony each from both MPN and control subjects were cultured in a monolayer in 96-well plates, stimulated with 100 ng/ml of LPS, and then tissue factor-dependent Xa generation estimated using a chromogenic substrate. There was some heterogeneity among different colonies, but as compared with the controls, MPN ECFCs had significantly elevated LPS-stimulated and tissue factor-dependent factor Xa generation (**Figure 3E**).

### ECFC transcriptomics highlight thrombo-inflammatory signature of MPN endothelium

To further characterize the role of endothelium in vascular inflammation associated with MPN, next, we carried out transcriptomic analysis of ECFCs patients with MPN compared with controls. RNA was extracted and bulk RNAseq performed on 5 control and 10 MPN ECFC colonies. A total of 289 genes were found to be differentially expressed in MPN ECFCs as compared with controls with an adjusted p-value of less than 0.05 and a fold-change cutoff of 1.5 (**Table S3**). Although some heterogeneity was notable, there was hierarchical clustering and distinct separation between MPN and control cohorts (**Figure 4A)**. The volcano plot highlights DEGs, with genes elevated in MPN shown in blue and those higher in controls in red (**Figure 4B**). Table 2 presents the top 10 most significant DEGs identified between MPN and control cohorts, with each row listing a gene, its encoded protein, and its biological function, while table 3 highlights DEGs specifically involved in thrombosis, angiogenesis and vascular inflammation.

**Figure 4.**
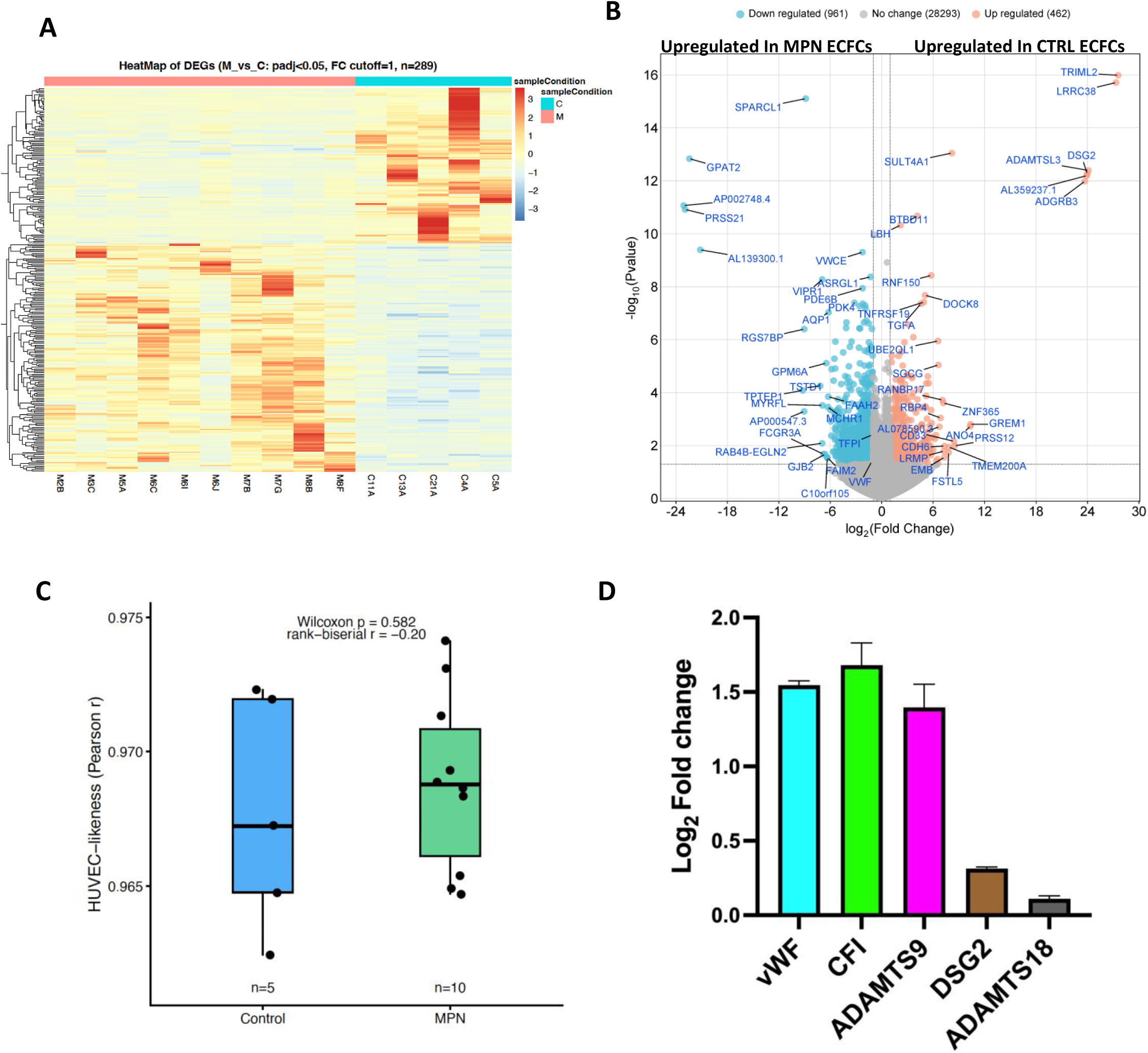
Transcriptomic profiling reveals endothelial dysfunction in MPN. (A) Z-score heatmap of differentially expressed genes (rows) between ECFCs from patients with MPN and controls (columns) (adjusted *p*-value<0.05; fold change (FC) >1.5). 289 DEGs were identified (red, upregulated; blue, downregulated. (B) Volcano plot of differentially expressed genes between MPN and control ECFCs (blue – higher in MPN; red, higher in controls; adjusted *p*-value<0.05; FC>1.5). (C) HUVEC-likeness (Pearson r) for each ECFC sample relative to the HUVEC reference; dots are samples, boxes show median/IQR, error bars are mean ± SD; two-sided Wilcoxon p and rank-biserial r shown. (D) Validation of selected differentially expressed genes (vWF, von Willebrand factor); CFI, complement factor I; ADAMTS9, A disintegrin and metalloproteinase with thrombospondin motifs 9; DS2, desmoglein 2; ADAMTS18, A disintegrin and metalloproteinase with thrombospondin motifs 18) identified by bulk RNA-seq in ECFCs from patients with MPN versus controls by qPCR; bars show log_2_ fold change (MPN vs control; deldel-Ct normalized to a housekeeping gene) plotted as mean ± SEM from independent biological samples.

**Table 2.** Top 10 DEGs identified in analysis of bulk RNAseq comparing MPN ECFCs with controls.

| *Gene* | *Protein* | *Function (Metascape; Database name:Biological Process (GO))* | *Log2 fold change* | *Adjusted P value* |
| --- | --- | --- | --- | --- |
| *TRIML2* | Tripartite motif family-like 2 | GO:0032526 response to retinoic acid;GO:0016567 protein ubiquitination;GO:0032446 protein modification by small protein conjugation | -27.57 | 1.85 E-12 |
| *LRRC38* | Leucine-rich repeat-containing protein 38 | GO:1903818 positive regulation of voltage-gated potassium channel activity;GO:1901018 positive regulation of potassium ion transmembrane transporter activity;GO:1901381 positive regulation of potassium ion transmembrane transport | -27.32 | 1.85 E-12 |
| *SPARCL1* | SPARC-like protein 1 (Hevin) | GO:0099560 synaptic membrane adhesion;GO:0098742 cell-cell adhesion via plasma-membrane adhesion molecules;GO:0050807 regulation of synapse organization | 8.86 | 4.99 E-12 |
| *SULT4A1* | Sulfotransferase family 4A member 1 | GO:0140059 dendrite arborization;GO:0051923 sulfation;GO:0140058 neuron projection arborization | -8.19 | 4.25 E-10 |
| *GPAT2* | Glycerol-3-phosphate acyltransferase 2, mitochondrial | GO:0006072 glycerol-3-phosphate metabolic process;GO:0052646 alditol phosphate metabolic process;GO:0016024 CDP-diacylglycerol biosynthetic process | 22.42 | 5.56 E-10 |
| *ADAMTSL3* | ADAMTS-like protein 3 | GO:0030198 extracellular matrix organization;GO:0043062 extracellular structure organization;GO:0045229 external encapsulating structure organization | -24.1 | 1.29 E-09 |
| *DSG2* | Desmoglein-2 | GO:0003165 Purkinje myocyte development;GO:0086073 bundle of His cell-Purkinje myocyte adhesion involved in cell communication;GO:0003164 His-Purkinje system development | -23.97 | 1.47 E-09 |
| *ADGRB3* | Adhesion G protein-coupled receptor B3 (BAI3) | GO:0061743 motor learning;GO:1905806 regulation of synapse pruning;GO:0016322 neuron remodeling | -23.67 | 2.20 E-09 |
| *PRSS21* | Testisin (Serine protease 21) | GO:0007283 spermatogenesis;GO:0048232 male gamete generation;GO:0007276 gamete generation | 22.94 | 2.09 E-08 |
| *BTBD11* | BTB/POZ domain-containing protein 11 | GO:0035640 exploration behavior;GO:0035249 synaptic transmission, glutamatergic;GO:0050821 protein stabilization | -4.16 | 3.38 E-08 |

**Table 3.**
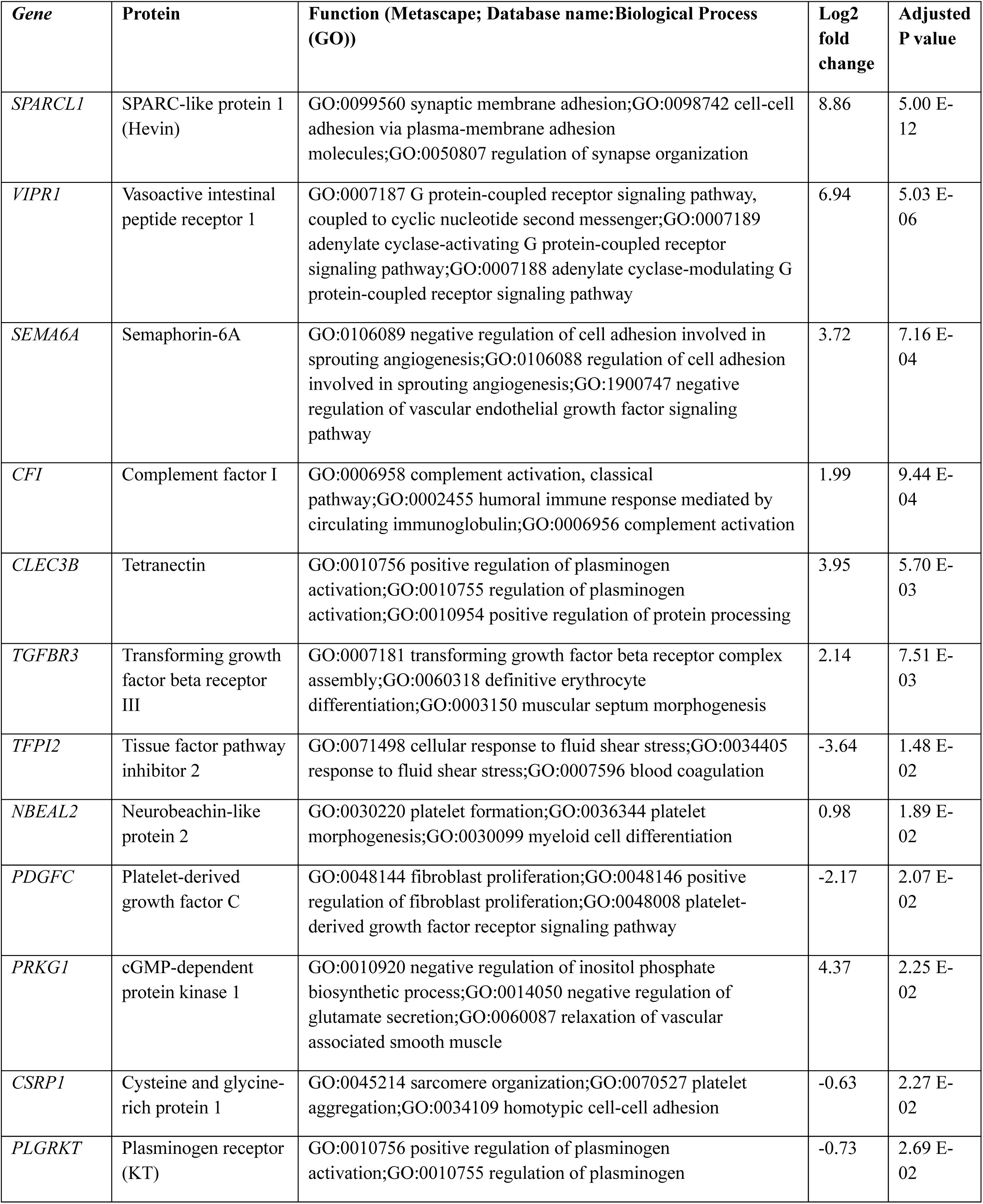
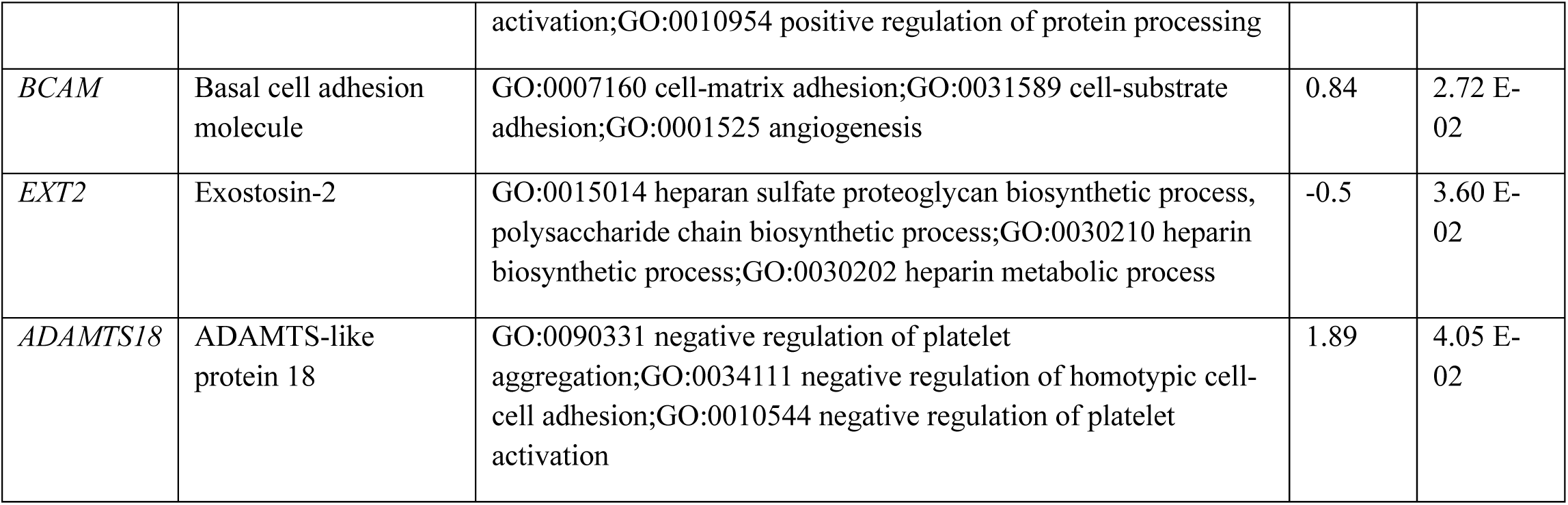
Top 15 DEGs with a functional role in vascular inflammation identified in the analysis of bulk RNAseq comparing MPN ECFCs with controls.

Given the origin and lineage of ECFCs remains unclear, to benchmark endothelial transcriptional identity, we compared bulk RNA-seq profiles from control and MPN ECFCs to an external primary endothelial reference (HUVEC). ECFC samples showed very high concordance with the HUVEC reference at the transcriptome-wide level, quantified as Pearson correlation to the HUVEC centroid expression profile (control mean r=0.9677, median r=0.9672; MPN mean r=0.9689, median r=0.9688), with no difference between control and MPN ECFCs (two-sided Wilcoxon p=0.594) (**Figure 4C and S4A-B)** This analysis indicate that ECFCs retain a robust endothelial transcriptional program and are strongly HUVEC-like, with no evidence for a disease-associated loss of endothelial identity in MPN-derived ECFCs.

To validate our transcriptomic data, we performed qPCR of a few DEGs identified above, particularly those with known functional roles in vascular inflammation. These included *vWF* and *DSG2* (high in MPN), as well as *CFI*, and *ADAMTS18* (low in MPN). **Figure 4D** shows the expression fold change of these genes in MPN as compared with controls. We also created an endothelial marker module score derived from canonical endothelial genes (e.g., *PECAM1*, *VWF*, *KDR/FLT1*, *EMCN*, *CLDN5* and related markers); this was found to be significantly higher in MPN ECFCs relative to controls (control mean −0.226, median −0.214; MPN mean 0.311, median 0.245; Wilcoxon p=0.019) (**Figure S4C-D**).

Next, we performed functional enrichment analysis of differentially expressed genes using the EnrichR gene set enrichment analysis suite.^28,30^ The top Gene Ontology (GO) biological process terms altered in MPN-derived ECFCs are shown in **Figure 5A** and **Table S4**, and include multiple pathways related to thrombosis and vascular inflammation, such as regulation of blood coagulation, negative regulation of platelet activation, regulation of secretion, and regulation of plasminogen activation. Additional enriched terms encompass processes involved in angiogenesis, proteoglycan and glycosaminoglycan biosynthesis, cellular energy metabolism, and intracellular homeostasis, indicating broad perturbation of endothelial function. To better visualize how these pathways interrelate, we generated a network plot of the most significantly enriched GO terms (**Figure 5B**), which reveals tightly connected clusters linking coagulation, platelet regulation, vascular remodeling, and metabolic stress.^31^ Highlighting representative terms from each cluster further underscores coordinated abnormalities in hemostatic balance, endothelial barrier and junctional regulation, extracellular matrix composition, and cellular stress responses in MPN ECFCs.

**Figure 5.**
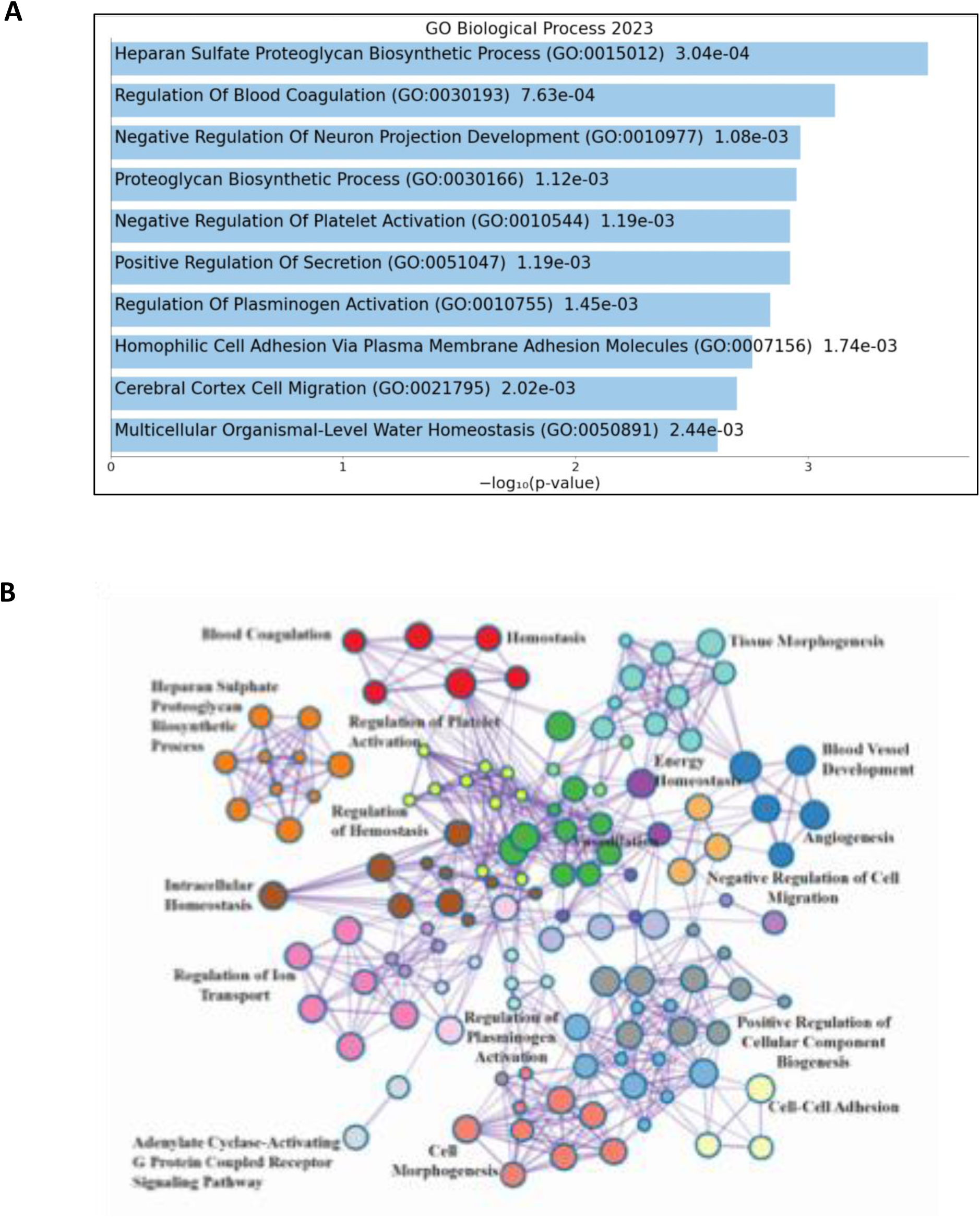
Enriched pathways highlight inflammatory and pro-thrombotic signatures in MPN endothelium. (A) Top 10 pathways (Gene Ontology) in MPN ECFC compared with controls derived from DEGs in figure 4A, along with corresponding *p*-values. (B) Network plot of enriched terms obtained from DEGs from MPN vs control ECFC comparison (figure 4A). Terms with a similarity of >0.3 are connected by edges. Each node represents and enriched term and is colored by its cluster ID. The most significant of the enriched terms are labeled in the network plot.

To confirm the analysis of our pathway analysis and to further determine whether MPN ECFCs exhibit an activated endothelial state, we next evaluated expression of many thrombo-inflammatory and cytokine-responsive endothelial transcripts. MPN ECFCs demonstrated significantly higher expression of *SELE* (E-selectin; control median 1.66 vs MPN median 3.24, p=0.0027) and *NFKBIA* (IκBα; control median 5.40 vs MPN median 6.08, p=0.0047), together with increased *VWF* expression (control median 10.46 vs MPN median 12.80, p=0.0127) (**Figure 6A**). In contrast, baseline *F3* (tissue factor) expression was low and not different between groups (p=0.859), suggesting that the prothrombotic/transcriptional shift in MPN ECFCs is driven primarily by selective activation programs (**Figure 6A**). Because canonical NF-κB signaling is controlled by inducible negative feedback circuits—most prominently through IκBα (*NFKBIA*) and A20 (*TNFAIP3*), which are NF-κB target genes that act to terminate and reset pathway activity—we quantified an NF-κB feedback module to assess baseline pathway tone/priming.^33^ Although selected NF-κB–responsive genes were increased in MPN ECFCs (notably *NFKBIA* and *SELE*), aggregated pathway module scores did not show a uniform baseline shift in NF-κB feedback or TNF/NF-κB activation programs (Wilcoxon p=0.513 and p=0.310, respectively) (**Figure 6 B and S5**). Overall, these findings support a model in which MPN ECFCs exhibit selective inflammatory priming.

**Figure 6.**
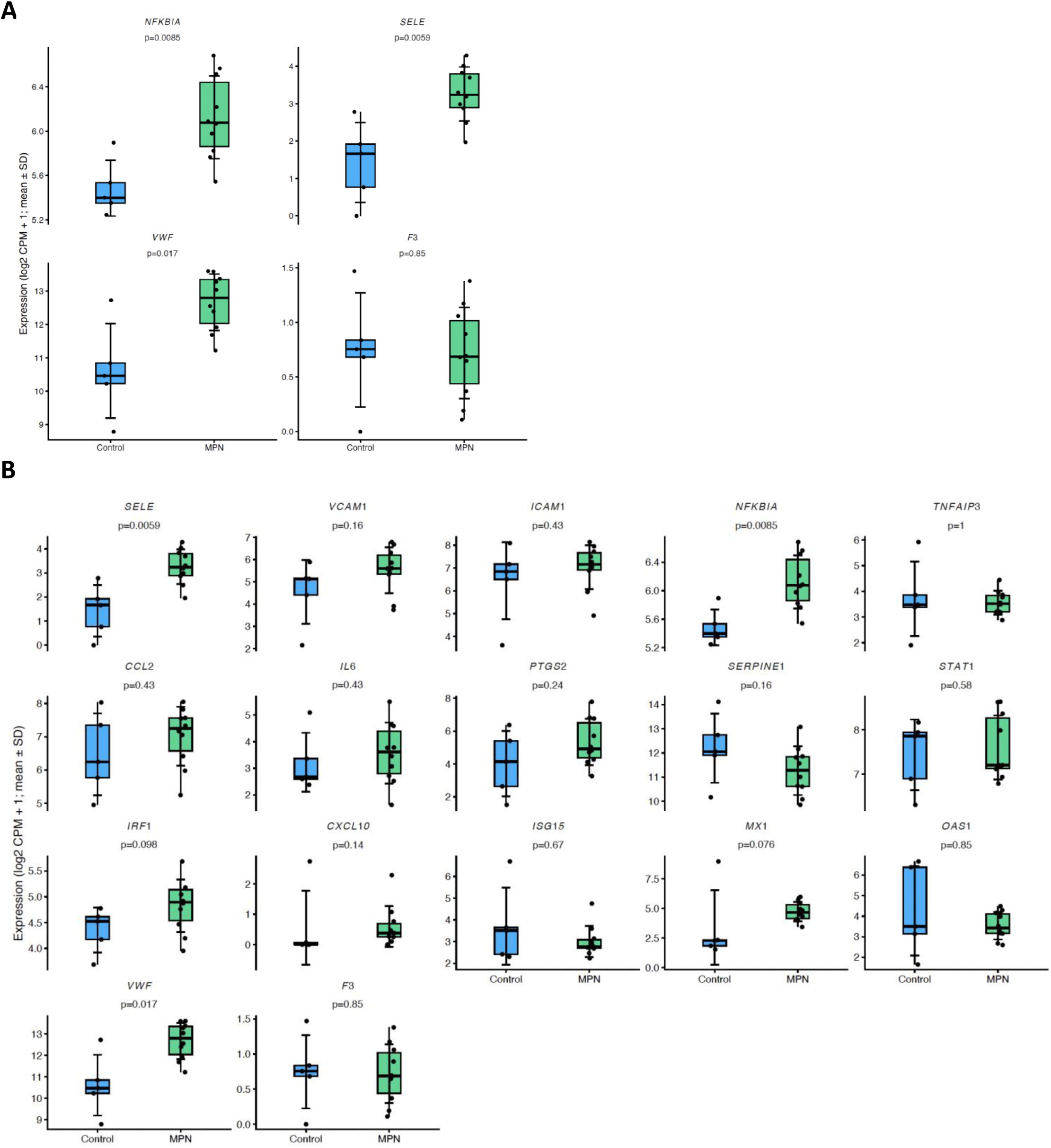
MPN ECFCs display thrombo-inflammatory endothelial activation with NF-kB/TNF priming. (A) Selected thrombo-inflammatory/endothelial genes (NFKBIA, SELE, VWF, F3) are shown as log2 CPM + 1 for Control vs MPN ECFCs; dots are individual samples, boxes show median/IQR with black mean ± SD error bars, and per-gene comparisons used two-sided Wilcoxon rank-sum testing (p and rank-biserial r displayed). (B) Expression of inflammatory and thrombo-inflammatory genes in Control and MPN ECFCs (log2 CPM + 1); dots represent individual samples, boxes show median/IQR with black mean ± SD error bars, and per-gene two-sided Wilcoxon rank-sum p-values are annotated in each facet.

Together, these pathway-level changes support a model in which MPN-derived endothelial cells adopt a prothrombotic and proinflammatory (“thrombo-inflammatory”) phenotype that may contribute to the heightened vascular risk observed in patients with MPN.

## Discussion

Thromboembolic events are the most common complication and a major cause of morbidity and mortality in patients with MPN.^6,34,35^ While peripheral blood leukocytes, particularly monocytes, and platelets have been extensively studied, the contribution of the vascular endothelium to the prothrombotic and inflammatory milieu in MPN is less well defined.^9,10,13^ Given the central role of endothelial cells in regulating hemostasis, leukocyte recruitment, and vascular tone, we set out to phenotypically and transcriptomically characterize primary endothelial cells from patients with *JAK2 V617F*–mutated MPN using ECFCs as a surrogate for vascular endothelium.

Our data support a model of endothelial activation and dysfunction in MPN. First, we observed a higher frequency and number of ECFC colonies in the MPN cohort compared with healthy controls, without any increase in proliferative capacity of MPN-derived ECFCs. This pattern is consistent with increased mobilization of endothelium or endothelial progenitors in the setting of chronic vascular inflammation, and aligns with prior reports of elevated CFU-ECs and ECFCs in MPN and other inflammatory conditions, despite some variability across studies.^12–16^

Second, MPN ECFCs exhibited features of a hyperreactive, prothrombotic endothelial phenotype. We previously demonstrated that plasma protein disulfide isomerase (PDI) is elevated in MPN and that endothelium, rather than platelets, is the principal source of circulating PDI, with higher levels associated with thrombotic events.^7^ In the current study, we extend these observations by showing increased expression and release of vWF and P-selectin by MPN ECFCs, key mediators of platelet adhesion, thrombosis, and leukocyte recruitment. Notably, we observed intra-subject heterogeneity in vWF and P-selectin expression across colonies, suggesting dynamic or locally regulated endothelial activation even within individual patients. Elevated endothelial vWF and P-selectin have been described in experimental models in which *JAK2 V617F* is overexpressed in endothelial cells, and in iPSC-derived ECs expressing mutant *JAK2*, all of which support a phenotype of endothelial hyperreactivity in MPN.^11,14–16,36–39^ Our findings in MPN patient-derived ECFCs, which do not themselves harbor *JAK2* mutations (see below), suggest that similar effector pathways can be engaged in a non-clonal endothelial compartment exposed to an MPN-associated inflammatory and procoagulant environment.

ECFCs have emerged as a useful model to study human endothelial biology in vivo–like conditions and have been employed to investigate vascular dysfunction in sickle cell disease, von Willebrand disease, rheumatologic diseases, and cardiovascular disorders.^20–22,24,25^ They exhibit many phenotypic and functional similarities to macrovascular endothelial cells such as HUVECs, including both phenotypic and transcriptomic profile. In MPN, both CFU-ECs (colony forming units – endothelial cells) and ECFCs have been studied, but the distinction between these populations is crucial when interpreting clonal status.

CFU-ECs are now recognized as hematopoietic in origin, express CD45, and may harbor *JAK2 V617F* and other myeloid driver mutations.^40–42^ In contrast, several studies, including ours, have shown that well-characterized, bona fide ECFCs from patients with *JAK2-*mutant MPN typically lack *JAK2 V617F* and other disease-defining mutations, supporting a non-clonal, non-hematopoietic endothelial lineage.^43,44^ Prior reports that had identified JAK2-mutant endothelial or endothelial-like cells in MPN and related conditions, did so often in settings where cell identity (CFU-EC vs ECFC vs endothelial progenitor cells) was less clearly defined or where different isolation strategies were used. Taken together, current data suggest that, at least in peripheral blood–derived ECFCs, endothelial dysfunction in MPN can occur in the absence of direct clonal involvement, likely driven instead by chronic exposure to inflammatory cytokines, activated blood cells, and altered hemostatic factors.

Moreover, because the developmental origin and in vivo counterpart of ECFCs remain incompletely defined, we benchmarked ECFC transcriptomes against an external, well-annotated endothelial reference (HUVEC RNA-seq processed through the same quantification pipeline). This analysis demonstrated a strong global transcriptional concordance between ECFCs and HUVECs, supporting that ECFC cultures capture a canonical endothelial gene-expression program. Importantly, although MPN ECFCs exhibited higher expression of select endothelial activation markers, these changes did not reduce overall “endothelialness” (HUVEC-likeness and endothelial identity scores), consistent with endothelial activation within an otherwise preserved endothelial state rather than lineage divergence or loss of endothelial identity.

Our transcriptomic analysis further supports the presence of endothelial dysfunction and thrombo-inflammation in MPN. We identified a set of differentially expressed genes in MPN ECFCs enriched for pathways related to angiogenesis, vascular permeability, extracellular matrix organization, and hemostatic regulation. Upregulation of *ADAMTS9*, *CFI*, *DSG2*, and *ADAMTS18*, among others, was confirmed by qPCR, implicating both complement and metalloprotease pathways in the prothrombotic phenotype. Functional enrichment analysis highlighted biological processes including regulation of blood coagulation, negative regulation of platelet activation, regulation of secretion, plasminogen activation, proteoglycan synthesis, and cellular homeostasis. Visualization of these terms in a network framework revealed interrelated clusters linking coagulation and platelet regulation to endothelial barrier function, extracellular matrix remodeling, and metabolic/stress responses. These findings align with spatial transcriptomic data from bone marrow vascular niches in MPN demonstrating enrichment of endothelial migration, sprouting, angiogenic, and inflammatory signaling programs (e.g., TNF/NF-κB, IL-1, and TGF-β pathways) in patients compared with healthy controls.^45^ They also complement prior work, that illustrated altered expression of ADAMTS17 and WNT pathway components in MPN ECFCs following co-culture with red cells.^37^

Collectively, these data suggest that the vascular endothelium in MPN, even when not clonally mutated, exists in a primed, thrombo-inflammatory state characterized by enhanced regenerative/activation capacity, increased expression and release of Weibel–Palade body contents, and broad transcriptional reprogramming of coagulation, adhesion, and inflammatory pathways.

This study has limitations. The sample size is modest and limits our ability to fully account for heterogeneity across MPN subtypes, mutational co-occurrences, and treatment exposures. ECFC culture is labor-intensive, requires several weeks for colony outgrowth, and may introduce selection biases toward more proliferative clones. Although ECFCs recapitulate many HUVEC-like endothelial features, their exact lineage and relationship to in vivo vascular beds remain incompletely defined, and findings may not fully generalize to all vascular territories relevant to MPN-associated thrombosis (e.g., splanchnic, cerebral, or microvascular beds). Our transcriptomic analyses were performed in bulk ECFC populations and do not capture single-cell heterogeneity or dynamic responses to inflammatory stimuli.

Despite these constraints, our work expands on a limited prior literature by providing an integrated phenotypic and transcriptomic characterization of patient-derived MPN ECFCs and placing endothelial dysfunction squarely within the broader framework of MPN thrombo-inflammation. Future studies in larger, mutation-stratified cohorts, incorporating single-cell and spatial transcriptomic approaches and functional co-culture models with clonal hematopoietic cells and platelets, will be essential to refine these observations and identify actionable targets. Nonetheless, our findings support the concept that the endothelium is not a passive bystander but an active participant in MPN pathobiology, contributing to a state of endothelial thrombo-inflammation that likely cooperates with clonal hematopoiesis to drive the heightened thrombotic risk characteristic of these disorders.

## Acknowledgements

This work was funded by National Institutes of Health (K08HL150246) to AVS. SASA received funding from the German Academic Exchange Service (DAAD) through the Biomedical Educational Program (BMEP) and the German Heart Foundation (DHS) through the Kaltenbach doctoral fellowship. We want to thank the Yale Keck Biotechnology Resource Laboratory and Yale Center for Genome Analysis (YCGA) for their help with RNA sequencing and analysis of data. We would also like to thank Prof. Dr. Michael Schuhmann, and the Graduate School of Life Sciences of the University of Wuerzburg for their support and supervision of the medical doctoral thesis of SASA.

## Author contributions

S.A.S.A. designed and performed experiments, analyzed data, and wrote the manuscript. A.D. and M.A. performed experiments and analyzed data. N.P. recruited study subjects and edited the manuscript. D.S. edited the manuscript. L.M.M designed experiments and edited the manuscript. A.V.S. designed and performed experiments, analyzed data, wrote and edited the manuscript.

## Conflict of interest

The authors declare no financial conflicts of interest.

## Notes

### Competing Interest Statement

The authors have declared no competing interest.

